# Combining NeuroPainting with transcriptomics reveals cell-type-specific morphological and molecular signatures of the 22q11.2 deletion

**DOI:** 10.1101/2024.11.16.623947

**Authors:** Matthew Tegtmeyer, Dhara Liyanage, Yu Han, Kathryn B. Hebert, Ruifan Pei, Gregory P. Way, Pearl V. Ryder, Derek Hawes, Callum Tromans-Coia, Beth A. Cimini, Anne E. Carpenter, Shantanu Singh, Ralda Nehme

## Abstract

Neuropsychiatric conditions pose substantial challenges for therapeutic development due to their complex and poorly understood underlying mechanisms. High-throughput, unbiased phenotypic assays present a promising path for advancing therapeutic discovery, especially within disease-relevant neural tissues. Here, we introduce NeuroPainting, a novel adaptation of the Cell Painting assay, optimized for high-dimensional morphological phenotyping of neural cell types, including neurons, neuronal progenitor cells, and astrocytes derived from human stem cells. Using NeuroPainting, we quantified cell structure and organelle behavior across various brain cell types, creating a public dataset of over 4,000 cellular traits. This extensive dataset not only sets a new benchmark for phenotypic screening in neuropsychiatric research but also serves as a gold standard for the research community, enabling comparisons and validation of results. We then applied NeuroPainting to identify morphological signatures associated with the 22q11.2 deletion, a major genetic risk factor for schizophrenia. We observed profound cell-type-specific effects of the 22q11.2 deletion, with significant alterations in mitochondrial structure, endoplasmic reticulum organization, and cytoskeletal dynamics, particularly in astrocytes. Transcriptomic analysis revealed reduced expression of cell adhesion genes in 22q11.2 deletion astrocytes, consistent with recent post-mortem findings. Integrating the RNA sequencing data and morphological profiles uncovered a novel biological link between altered expression of specific cell adhesion molecules and observed changes in mitochondrial morphology in 22q11.2 deletion astrocytes. These findings underscore the power of combined phenomic and transcriptomic analyses to reveal mechanistic insights associated with human genetic variants of neuropsychiatric conditions.

## Introduction

Severe mental illnesses represent a significant global health crisis. Individuals suffering from mental illness face debilitating symptoms, including dramatic mood swings, persistent hallucinations, and increased risk of substance abuse and suicidality, contributing to severe personal and societal consequences such as unemployment, homelessness, and incarceration ^1,2^. Current therapeutic approaches, largely unchanged for decades, rely on a combination of psychotherapy and pharmacotherapy. However, these treatments do not adequately address the needs of all patients, with more than half of people with schizophrenia and bipolar disorder continuing to experience debilitating functional impairments ^3–6^. This inadequacy underscores a critical gap in our understanding of the biological mechanisms of these conditions, hindered further by the slow pace of innovation in psychiatric therapeutic development. The stalled advancement in therapeutic options is partly due to the complex nature of these disorders, which are influenced by a myriad of genetic and environmental factors ^7^. There is a need for novel, robust phenotyping approaches to increase our ability to extract meaningful biological insights associated with these conditions.

Chromosome 22q harbors the recurrent 22q11.2 microdeletion (22q11.2del), a well-studied copy number variant (CNV) that leads to the 22q11.2 deletion syndrome (also known as DiGeorge Syndrome), a multisystem disorder characterized by a range of neuropsychiatric, cardiac, and immune phenotypes ^8^. Notably, 22q11.2del is the genetic factor most strongly associated with schizophrenia at the population level ^9^. The cardiac effects of this deletion have been linked to the *TBX1* gene within the canonical deletion interval ^10^. However, the neuropsychiatric impacts remain largely unexplained by candidate genes within the deleted region, suggesting the possibility of non-canonical mechanisms underlying these phenotypes ^10–13^. Additionally, like all other chromosomes, 22q contains thousands of common SNPs that contribute additively to human complex trait variation, including traits associated with the 22q11.2 deletion ^14^. Given the immense clinical heterogeneity of 22q11.2del ^15^, robust, multimodal phenotyping is crucial to uncover the molecular mechanisms driving this disorder. Such an approach will help identify affected pathways and assess the mutation’s impact across different cellular contexts, offering insights into its diverse clinical manifestations, with potential implications for schizophrenia more broadly.

Phenotypic assays such as Cell Painting ^16^ that capture high-throughput, unbiased cellular morphological measurements are appealing because they do not require selecting a specific target nor even a specific phenotype in advance, both of which can be flawed if based on incomplete mechanistic understanding. Cell Painting, and its adaptations, have been used in various applications to understand gene function, uncover cell-type specific biology, and build large reference datasets for the scientific community ^17–21^. Cell morphology profiling offers a significant advantage for functional genomics studies over methods like gene expression, as it is more cost-effective and easily scalable for both bulk and single-cell level analyses of mRNA. Cell morphology profiling also has the advantage of capturing downstream phenotypic outcomes, whereas mRNA levels reflect the potential for protein expression and may show only partial correlation with protein levels ^22^. Cell morphology profiling therefore allows for a more direct assessment of functional cellular changes in response to genetic or environmental perturbations.

Recent studies indicate that disease-associated cellular phenotypes can be identified directly from patient samples in an unbiased manner, rather than relying on hypothesized phenotypes. Specifically, morphology phenotypes have been discovered for neurological and neuropsychiatric conditions by image-based profiling of primary skin fibroblasts. For example, mitochondrial morphology was disrupted in a study of 41 patients with mid-stage idiopathic Parkinson’s disease compared to controls ^23^. Similarly, a deep learning-based analysis identified distinct morphological phenotypes in 46 Parkinson’s patients—32 sporadic and 14 with specific mutations—compared to healthy controls ^24^. Machine learning also differentiated images of 12 spinal muscular atrophy patients from 12 healthy controls, though technical confounders could not be ruled out ^25^. Wali et al. successfully used morphological profiling, including DNA, mitochondria, and acetylated α-tubulin stains, to distinguish 15 patients with hereditary spastic paraplegia and matched controls ^26^. While associating a cell phenotype with a condition does not prove causation, using patient-derived cellular phenotypes provides a strong link to therapeutic hypotheses: reversing an unhealthy cellular phenotype through chemical modulation *in vivo* may be effective at reversing behavioral deficits.

A strength of studying primary cells, such as skin fibroblasts, is their ease of collection, allowing studies to reach sufficient power to elucidate cellular phenotypes. The use of primary somatic cells also presents a limitation. For some conditions, including psychiatric disorders, it is important to consider phenotypes in the appropriate tissues, as disease-associated genetic variation is often enriched in tissue-specific genes ^27,28^. Wali et al. recognized this and extended their initial work on fibroblasts to patient-derived induced pluripotent stem cells (iPSCs) to validate phenotypes in neural cells ^29^. While this shift to iPSCs allowed for the examination of more disease-relevant cell types, it also significantly reduced the study’s scale, limiting phenotyping to only six cell lines (three healthy/three cases) and focusing on a few cellular characteristics like mitochondria and neurite length. Although this approach was useful for identifying strong phenotypes from highly pathogenic mutations, it underscores the need for methods capable of detecting more subtle phenotypes associated with milder genetic mutations that may present with significant clinical heterogeneity^30,31^. To achieve this, it will be necessary to extend these studies to a larger number of cell lines and capture a broader range of cellular features. State-of-the-art microscopy approaches like Cell Painting, which can measure thousands of morphological features ^16,17^, offer a promising solution.

Here, we established NeuroPainting, a high-dimensional phenotyping assay adapted from Cell Painting, for morphological discovery in neural cell types derived from human iPSCs. This robust and scalable system enabled us to examine cells from many donors simultaneously, increasing the likelihood of capturing individual variation known to exist at the genetic and clinical levels in psychiatric conditions. We discuss the optimization and development of the NeuroPainting assay and analytical pipelines and quantify cellular structure and organelles across different neuropsychiatric relevant iPSC-based neural cell types, including neurons and astrocytes. We further use NeuroPainting to discover morphological phenotypes associated with the 22q11.2 deletion. We show that NeuroPainting can quantify differences among diverse cell types and demonstrate cell-type specific morphological phenotypes linked to the 22q11.2 deletion, highlighting the utility of NeuroPainting for uncovering new biology linked to genetic variants implicated in psychiatric disease. Lastly, we integrated RNA sequencing data with our NeuroPainting morphological data, pinpointing specific cell adhesion genes that may underlie morphological changes in astrocytes with 22q11.2 deletion and suggesting new avenues for biological characterization and potential therapeutic strategies.

## Results

### NeuroPainting for high-dimensional morphological phenotyping of neural cell types at scale

We developed “NeuroPainting” by adapting the six original dyes of the Cell Painting assay to permit robust and scalable analyses of iPSC-derived neural cell types (**Fig. 1a**). We generated NeuroPainting profiles from stem cells, neuronal progenitor cells, neurons, and astrocytes from 44 cell lines, 22 of which harbor a 22q11.2 deletion and 22 neuro-typical controls with matched age and ancestry, which we previously characterized ^13^ (**Fig. 1b, Table S1**). For each cell type, all lines were differentiated in concert using methods we previously described ^32–34^ before being plated in 384-well microplates for staining and imaging. To reduce technical variation known to impact microscopy-based assays ^24^, we randomized plate maps to ensure equal distribution of genotypes and cell lines across the plates. We plated cells at specific densities and matured them to a particular time point determined by preliminary experiments (**Methods**) before fixing and staining. Cells were then imaged at 20X using a Perkin Elmer Phenix high-content imaging system.

**Figure 1.**
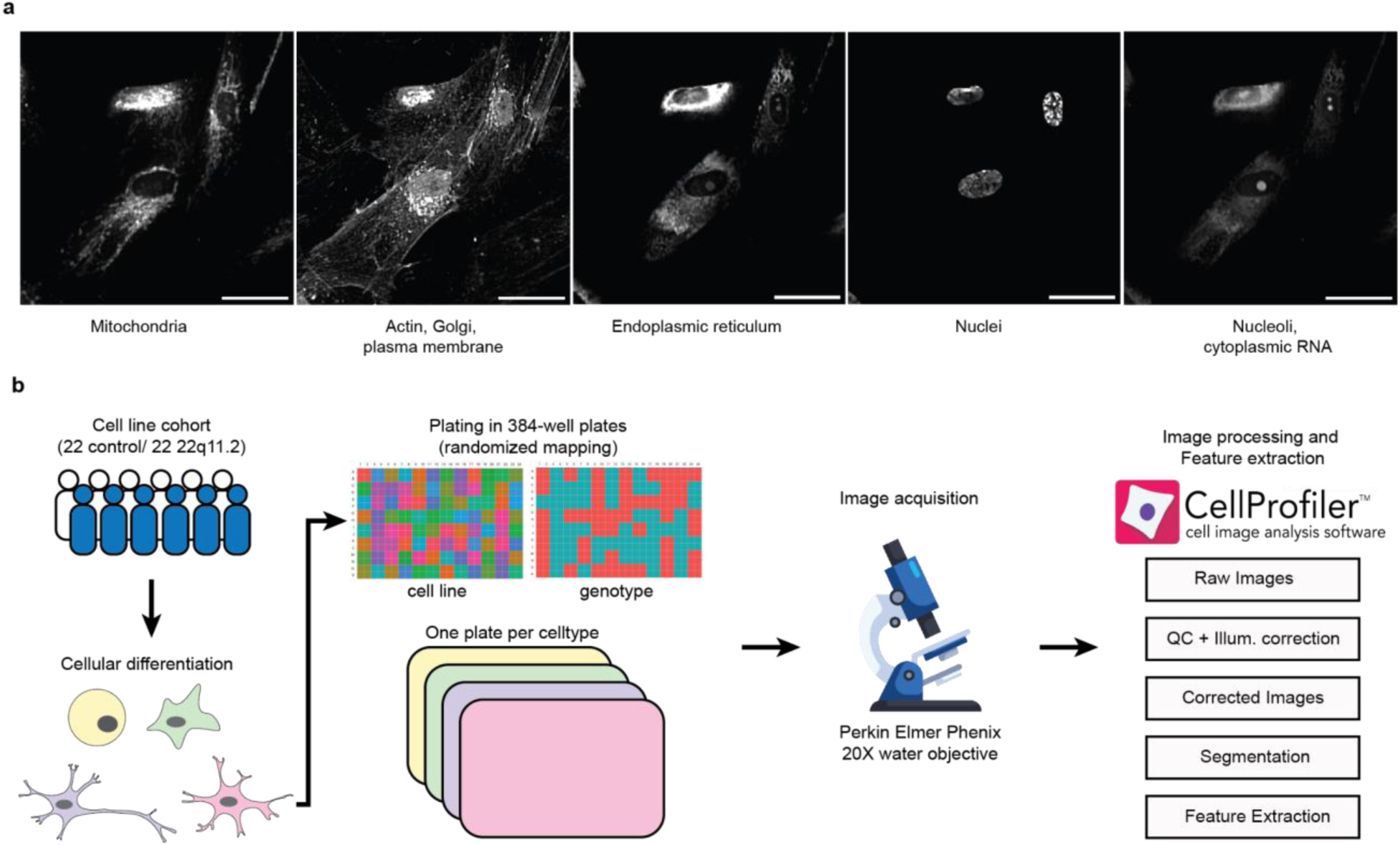
Study overview. **a.** Representative images of astrocytes stained with the Cell Painting dyes. Scale bar = 50 uM. **b**. Schematic of strategy for capturing NeuroPainting high-dimensional image-based profiles from a large cohort of iPSCs.

We created an image analysis and feature extraction pipeline specific to neural cell types in the freely available software, CellProfiler. We chose standard methods for feature extraction (as opposed to deep learning based methods) to improve interpretation of morphological phenotypes - well-understood features of morphology include size, shape, intensity, and texture of various stains which resulted in 4,104 different cellular traits ^35^. We grouped these traits based on each cellular compartment (Cells, Cytoplasm, Nuclei) and seven measurement categories: AreaShape, Granularity, Intensity, Radial Distribution, Correlation, ObjectSkeleton, and Texture (**Table S2**). Measurements in an additional “Other” category were excluded from analysis, because it includes some non-biological features such as location within the well. All analytical pipelines are freely available via our GitHub repository (see **Methods**). We refer to the collection of single-cell features measured by these pipelines as NeuroPainting profiles. We then mean-averaged single-cell profiles to create a profile for each well. These well-level average profiles underwent a four-step preprocessing pipeline described in ^36^, yielding 723 unique features (**Table S3**): (1) removal of low-variance features (coefficient of variation <1e-3), (2) robust standardization using median absolute deviation, (3) rank-based inverse normal transformation, and (4) correlation-based feature selection to eliminate redundant features with correlation >0.9.

### Morphology-Based Classification of Neural Cell Types

By eye, the NeuroPainted cell types displayed markedly different morphology (**Fig. 2a**).

**Figure 2.**
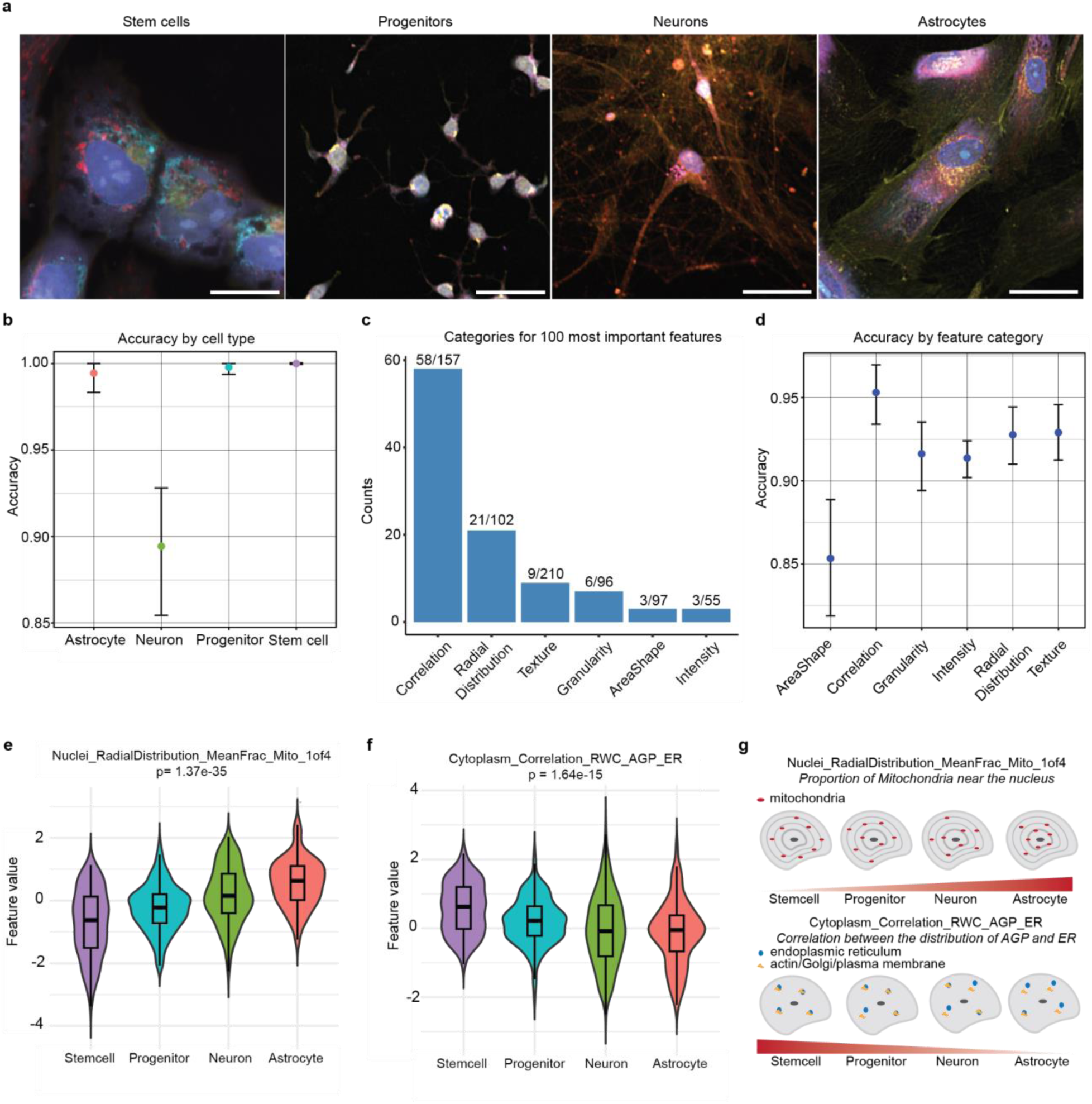
Cell type specific morphological signatures. **a.** Representative images of cell types included in this study. Scale bar = 50 uM. **b.** Random Forest classification accuracy across cell types. Data are presented as dot plots, where each point represents the mean accuracy across cross-validation folds for each feature category. Confidence intervals are computed using bootstrapping, and the error bars indicate the 95% confidence intervals around the mean. The size of the dots reflects the overall accuracy, and the error bars span the range of bootstrapped estimates of the mean. **c.** Barplot of feature categories for 100 most important features based on our Random Forest classifier. The fractions of the features marked as important are indicated for each category. **d.** Random Forest classification accuracy for cell types across feature categories, presented similarly to Figure 2b. **e.** Violin plot for *Nuclei_RadialDistribution_MeanFrac_Mito_1of4* across cell types (p = 1.37e-35, ANOVA). Data are presented in a Turkey-style boxplot with the median (Q2) and the first and the second quartiles (Q2, Q3) and error bars defined by the last data point within ±1.5-times the interquartile range. **f.** Violin plot for *Cytoplasm_Correlation_RWC_AGP_ER* across cell types (p = 1.64e-15, ANOVA), presented similar to Fig 2e. **g.** Schematic representation of two distinguishing cell morphology characteristics across the four cell types.

While, using our approach, each cell type is generated separately, the ability to classify cell types based on morphological profiles would be extremely valuable for examining cultures or tissues with mixed cell types, and for studying the effect of mutations, or genetic or pharmacological perturbations on cell states and differentiation potential and trajectories. To explore whether we could use NeuroPainting to classify cell types, we compared profiles across stem cells, neuronal progenitors, neurons, and astrocytes from the neuro-typical controls. Although the differentiation protocol for each cell type was distinct, each imaging batch contained all donors of a given cell type. This setup facilitated robust cross-donor comparisons—our primary goal—but we acknowledge that it could introduce batch effects across cell types.

We trained a Random Forest classifier to distinguish among the four cell types based on 723 features across 788 samples. The model achieved 97.41% overall accuracy (95% CI: 92.63%–99.46%) with a Kappa of 0.962, indicating strong agreement between predicted and true cell types. Neurons were classified perfectly (Sensitivity = 1.00), while astrocytes (0.92), progenitors (0.96), and stem cells (0.96) had slightly lower sensitivities. Positive predictive values (PPV) were similarly high, though neurons had a slightly lower PPV (0.95), while the other cell types achieved perfect values. Balanced accuracy ranged from 95.83% to 98.08%, confirming the model’s robust performance (**Fig. 2b**).Thus, the classifier confirmed that NeuroPainting accurately captures meaningful, biologically relevant differences in cell morphology.

To identify the most important features, we ranked their contributions to the model’s performance. Correlation and RadialDistribution features dominated the top 100, comprising 80% of the most informative features for classifying cell types (**Fig. 2c**). This result was unexpected, as we had anticipated a more prominent role for AreaShape features, given the observable differences in cell shape.

To further assess the contribution of each feature category, we retrained our classifier using only one of the six feature types at a time (AreaShape, Correlation, Granularity, Intensity, RadialDistribution, and Texture). The Correlation category achieved the highest performance (accuracy = 95.31%, Kappa = 0.93), followed by Texture (accuracy = 92.90%, Kappa = 0.89) and RadialDistribution (accuracy = 92.76%, Kappa = 0.89). AreaShape had the lowest accuracy (85.34%, Kappa = 0.78), while Granularity and Intensity both showed moderate performance, with accuracies around 91% and Kappa values of 0.87. These findings underscore the importance of Correlation and Texture features, while AreaShape features, although less powerful individually, still contribute meaningfully to distinguishing cell types (**Fig. 2d**).

From the top of our feature importance analysis, we highlighted two key features: Nuclei_RadialDistribution_MeanFrac_Mito_1of4, which relates to mitochondrial distribution around the nucleus, and Cytoplasm_correlation_RWC_AGP_ER, which measures the relative weighted correlation between actin/Golgi/plasma membrane and the endoplasmic reticulum in the cytoplasm. Both features exhibited statistically significant differences across cell types (**Fig. 2e**, f; ANOVA, p = 1.37e-35, p = 1.64e-15). We have included schematics to illustrate these features and their behavior in different cellular contexts (**Fig. 2g**).

In conclusion, our results suggest that NeuroPainting provides a robust and biologically meaningful approach to classifying neural cell types. Beyond classification, it offers a nuanced understanding of how cell morphology diverges across lineages, complementing gene expression data. This capability opens the door to using NeuroPainting as a powerful readout in studies exploring functional consequences of genetic variation and treatment responses in neurobiology.

### Cell-type specific morphological signatures in neural cell types from individuals with 22q11.2 deletion syndrome

The 22q11.2 deletion is the most common chromosomal deletion, associated with various diseases including congenital heart defects, autoimmune disorders, and neuropsychiatric conditions such as schizophrenia ^8^. While prior research has shown that this deletion alters gene expression and synaptic function in specific cell types ^13^, the underlying mechanisms driving its broad phenotypic effects remain poorly understood. This highlights the need for high-throughput phenotyping to uncover new insights. To address this, we employed NeuroPainting to investigate whether morphological signatures could differentiate control and deletion samples across neural cell types, potentially offering new avenues for drug screening.

We generated NeuroPainting profiles from neuronal progenitor cells, neurons, and astrocytes derived from iPSCs of individuals with 22q11.2 deletion syndrome and matched controls. Correlation matrices revealed distinct clustering of control and deletion samples, indicating clear morphological differences among these groups across cell types (**Fig. 3a, Fig. S1a**). Pairwise Pearson correlations within and between conditions were calculated for each cell type. The mean correlation for control-control pairs was r = 0.17, 0.03, 0.007, and 0.14 for stem cells, progenitors, neurons, and astrocytes, respectively. These values dropped substantially in control-deletion comparisons, with r = -0.11, -0.02, -0.003, and -0.10. A two-sample t-test confirmed that these distributions were significantly different for stem cells, progenitors, and astrocytes, but not for neurons (t-statistic = 96.09, p = 4.68e-53; t-statistic = 14.70, p = 9.279608e-49; t-statistic = 7.28, p = 3.165349e-13; t-statistic = 33.86, p = 3.948375e-232) (**Fig. 3b**). This demonstrates that the 22q11.2 deletion induces distinct morphological changes in these cell types.

**Figure 3.**
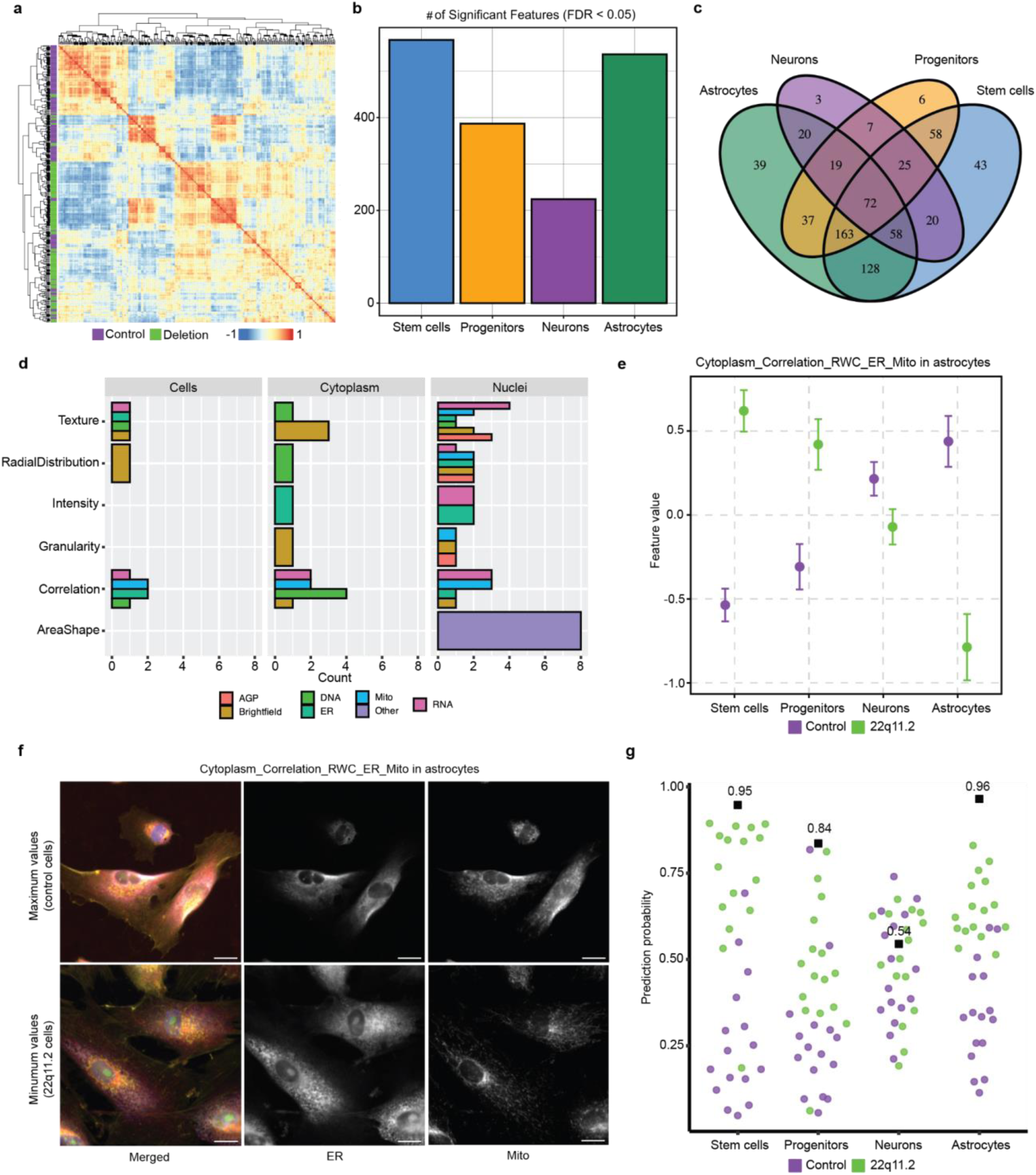
Genotype-specific morphological signatures in 22q11.2 deletion syndrome. **a.** Correlation matrix of NeuroPainting profiles for stem cells from control and 22q11.2 deletion cell lines. The green and purple notation on the left of the plot indicates control versus deletion donors. **c**. Venn diagram showing the number of significant features that overlap across the different cell types. **d.** Barplot of feature characteristics for 72 overlapping significant features. **e**. Dotplot of feature values for *Cytoplasm_Correlation_RWC_ER_Mito* across each cell type. Each data point represents a specific gene-feature pair, with different colors used to distinguish pairs. Error bars represent the standard error of the mean for both expression (horizontal) and morphology (vertical). **f**. Representative images for min/max values for *Cytoplasm_Correlation_RWC_ER_Mito* in control (top) and 22q11.2 (bottom) astrocytes. Images were selected based on highest and lowest well-average feature value. **g**. Scatter plot for AUC scores from donor-level NeuroPainting profiles across cell types, colored by genotype.

Next, we performed a Wilcoxon Rank-Sum test to identify specific morphological features significantly impacted by the deletion. We identified 567, 387, 224, and 536 significant features in stem cells, progenitors, neurons, and astrocytes, respectively that passed multiple hypothesis correction (**Fig. 3c**, **Tables S4-S7**, Wilcoxon, FDR < 0.05). Among these, 72 significant features overlapped across each cell type, suggesting some shared effects of the 22q11.2 deletion in different cellular contexts (**Fig. 3d**). Many feature categories, including texture, radial distribution, and correlation across several organelles were altered by the mutation in all cell types (**Fig. 3e**). The most prominent impact of the 22q11.2 deletion across cell types was associated with the AreaShape of the Nuclei (**Fig. 3e**).

The effect of the 22q11.2 deletion on *Cytoplasm_Correlation_RWC_ER_Mito* showed an interesting response: higher values for the deletion in stem cells and progenitors, roughly equivalent values for neurons and lower values for astrocytes (**Fig. 3f**); this cell type-specific behavior parallels previous findings from gene expression analyses ^13^. This feature describes the spatial relationship between the endoplasmic reticulum (ER) and mitochondria in the cytoplasm, implying that the deletion disrupts the normal ER-mitochondria interaction (**Fig. 3g**). The majority of significant features did not overlap, pointing to cell-type specific effects of the 22q11.2 deletion on cell morphology (**Fig. S1b**). These mirrored earlier observations by our group when examining gene expression patterns in these same cell lines ^13^.

In stem cells, numerous texture-based features for AGP, RNA, and mitochondria across all cellular compartments were identified, indicating potential organelle dysfunction (**Fig. 3h,** p=2.69e-08). Neuronal progenitor cells carrying a 22q11.2 deletion showed altered cell size (**Fig. 3i,** p=0.000268), consistent with neuroimaging studies linking the mutation to changes in brain size ^37,38^. Neurons with the 22q11.2 deletion exhibited asymmetry of their cell bodies when compared to control cells (**Fig. 3j,** p = 0.0239). In astrocytes, the deletion resulted in altered radial distributions of the ER, AGP, and mitochondria, potentially impacting cell shape, motility, and mitochondrial dynamics (**Fig. 3k,** p = 5.9e-6).

We next sought to determine whether the observed differences in NeuroPainting signatures could be leveraged to classify samples by genotype. For each donor and cell type, we averaged individual morphological features and applied a Random Forest classifier. The classifier achieved high accuracy across most cell types, with balanced accuracies of 0.91 for stem cells (p < 0.0001), 0.96 for astrocytes (p < 0.0001), and 0.84 for progenitors (p = 0.04). However, classification accuracy was notably lower in neurons (0.56, p = 0.30), consistent with fewer differentially significant features in this cell type. The area under the curve (AUC) scores further supported these findings, indicating strong discriminatory power in stem cells (AUC = 0.95) and astrocytes (AUC = 0.96), moderate performance in progenitors (AUC = 0.84), and poor performance in neurons (AUC = 0.54) (**Fig. 3g**). These results suggest that NeuroPainting signatures reliably capture genotype-specific morphological differences, with limited retrievability in neurons.

These findings highlighted widespread disruptions in cellular organization across cell types associated with the 22q11.2 deletion.

### 22q11.2 deletion regulates expression of cell-adhesion molecules in astrocytes

We next further investigated the impact of the 22q11.2 deletion in astrocytes, which displayed nearly 408 significantly altered morphological features. Given increasing evidence of the critical role that glial cells, particularly astrocytes, play in psychiatric conditions through astrocyte-neuron interactions ^34,39–41^, we aimed to understand how this deletion affects astrocyte function.

We generated genome-wide gene expression data from a subset of our cell lines (n=12, 6 control and 6 22q11.2) using a pooled (cell village) approach (Mitchell et al., 2020; Wells et al., 2023). Uniform Manifold Approximation and Projection (UMAP) showed high homogeneity across individual cell lines, with no cell line clustering distinctly from others using Louvain unsupervised clustering (**Fig. 4a**). Nor did we observe dramatic clustering based on genotype, indicating that the presence of the deletion did not impair differentiation capacity broadly (**Fig. 4b**). The fact that overall expression profiles were not dramatically different suggested that a consistent change in expression of individual genes might be informative. As expected, we observed a consistent decrease in the expression of genes located in the Chr22q11.2 interval (**Fig. S2**).

**Figure 4.**
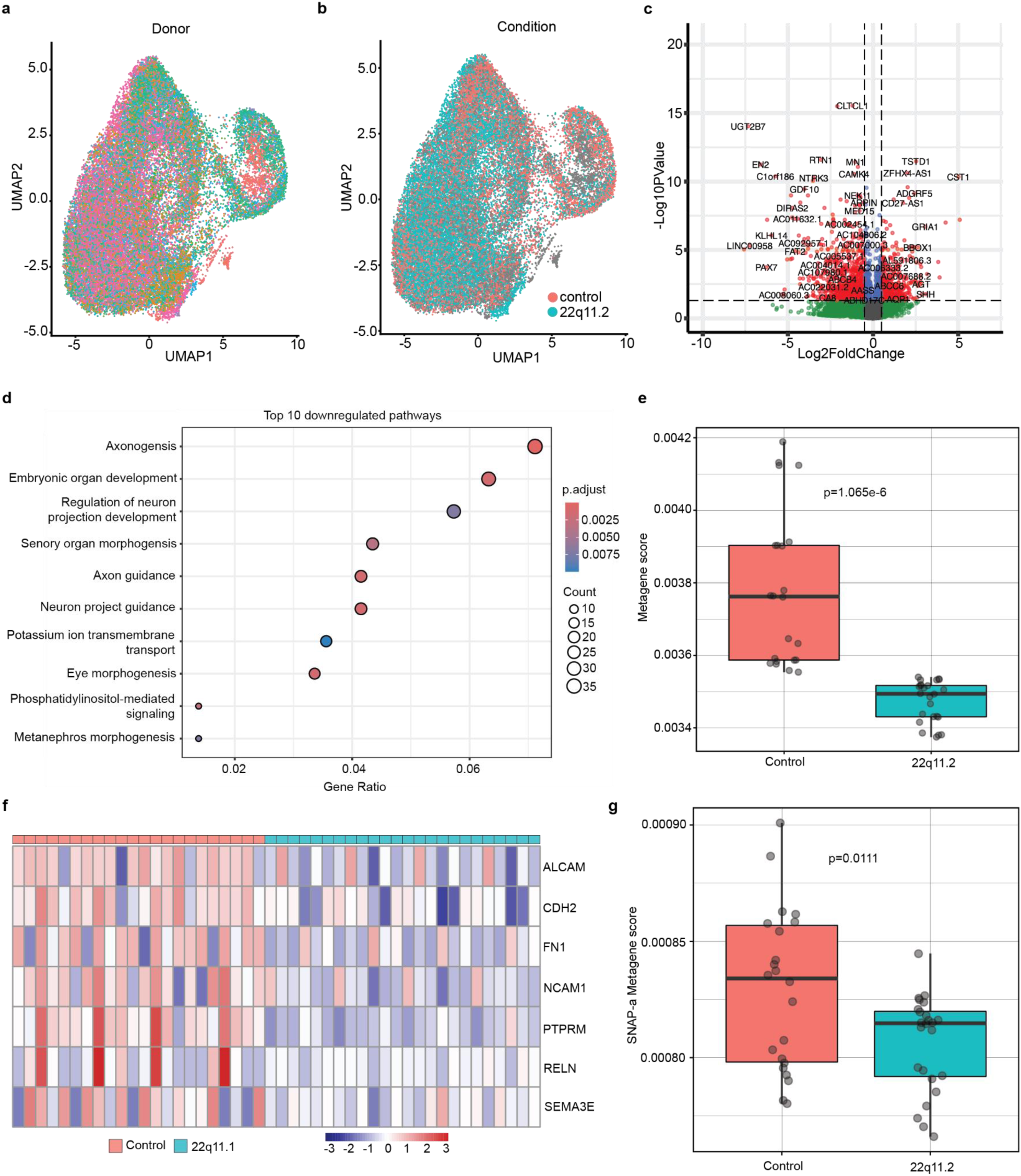
Dysregulation of cell-adhesion mRNA in 22q11.2 astrocytes. **a.** UMAP projection of single cell RNA-sequencing (scRNAseq) data from astrocytes colored by donor. **b**. UMAP projection of astrocytes scRNAseq data colored by genotype. **c**. Volcano plot for differentially expressed genes (Log2FC cutoff = 0.05, padj > 0.05). **d**. Gene set enrichment for downregulated genes. **e.** Metagene score analysis for genes in downregulated pathways. Data are presented like 2e. **f**. Heatmap of logCPM expression for cell-adhesion genes. **g**. Metagene score analysis for top 25 SNAP-a genes (*NRXN1, SLC1A2, RNF219-AS1, NTM, ZNF98, GPC5, GRM3, HPSE2, NKAIN3, SLC4A4, CTNND2, NCKAP5, SGCD, LSAMP, GPM6A, LRRC4C, LRRTM4, EPHB1, PREX2, RORA, TMEM108, ARHGAP24, SYNE1, TENM2, AC091826.2*). Data are presented like 2e.

We measured differentially expressed genes (DEGS) using DESeq2 by generating pseudo-bulked expression profiles for each donor across sequencing replicates (**Methods**). We identified 1358 DEGS based on our threshold (**Fig. 4c**, **Table S8**, Log2FC > 0.5, padj < 0.05). We next performed a gene-set enrichment analysis on downregulated genes in 22q11.2 deletion astrocytes. Several pathways related to neuronal development and axon generation and guidance were among the most downregulated (**Fig. 4d**). We observed a strong decrease in the expression of genes in these pathways by applying a metagene approach (Two-sample T-test, t = 6.5022, df = 23.712, p = 1.065e-06). Notably, many of these genes (*ALCAM, CDH2, FN1, NCAM1, PTPRM, RELN, SEMA3E*) behave as cell-adhesion molecules and have been linked to schizophrenia and other psychiatric conditions (**Fig. 4f**) ^42–48^. Recent studies have shown that disruption of cell-adhesion gene expression in astrocytes is associated with schizophrenia, and these genes harbor a high degree of genetic risk for psychiatric outcomes ^39,49^. To assess whether the impact of the 22q11.2 deletion on astrocyte gene expression was consistent with emerging evidence from findings in brain tissue from persons with idiopathic schizophrenia, we generated a metagene score based on the top gene loadings in astrocytes from the synaptic-neuron astrocyte program (SNAP) identified in Ling et al. We observed a significant decrease in the expression of these genes in 22q11.2 astrocytes compared to control cells (**Fig. 4g**, two-sample T-test, t = 2.6843, df = 33.663, p = 0.01119).

These results suggested that the 22q11.2 deletion significantly impacts astrocyte gene expression patterns relevant to psychiatric conditions.

### Gene-morphology correlations in 22q11.2 deletion highlight changes in mitochondrial features

We next sought to explore the relationship between these changes in gene expression and changes in cell morphology to nominate genes which may play a role in our 22q11.2 NeuroPainting signatures. To do this, we generated donor-level (n=12, control=6/deletion=6) gaussianized profiles for both RNA-seq and our morphological profiles in astrocytes. For our analyses, we focused on all DEGs (1358) and all significant morphological features (536) from our earlier results.

A Canonical Correlation Analysis (CCA) identified strong associations between gene expression patterns and morphological features, with the first canonical variable (CV1) effectively separating samples by condition, indicating that the relationships between gene expression and morphology were significantly altered in the deletion samples compared to controls (**Fig. 5a**). To further understand these relationships, we examine the top 100 genes and top 100 morphological features based on their loadings in the first canonical variable (CV1).

**Figure 5.**
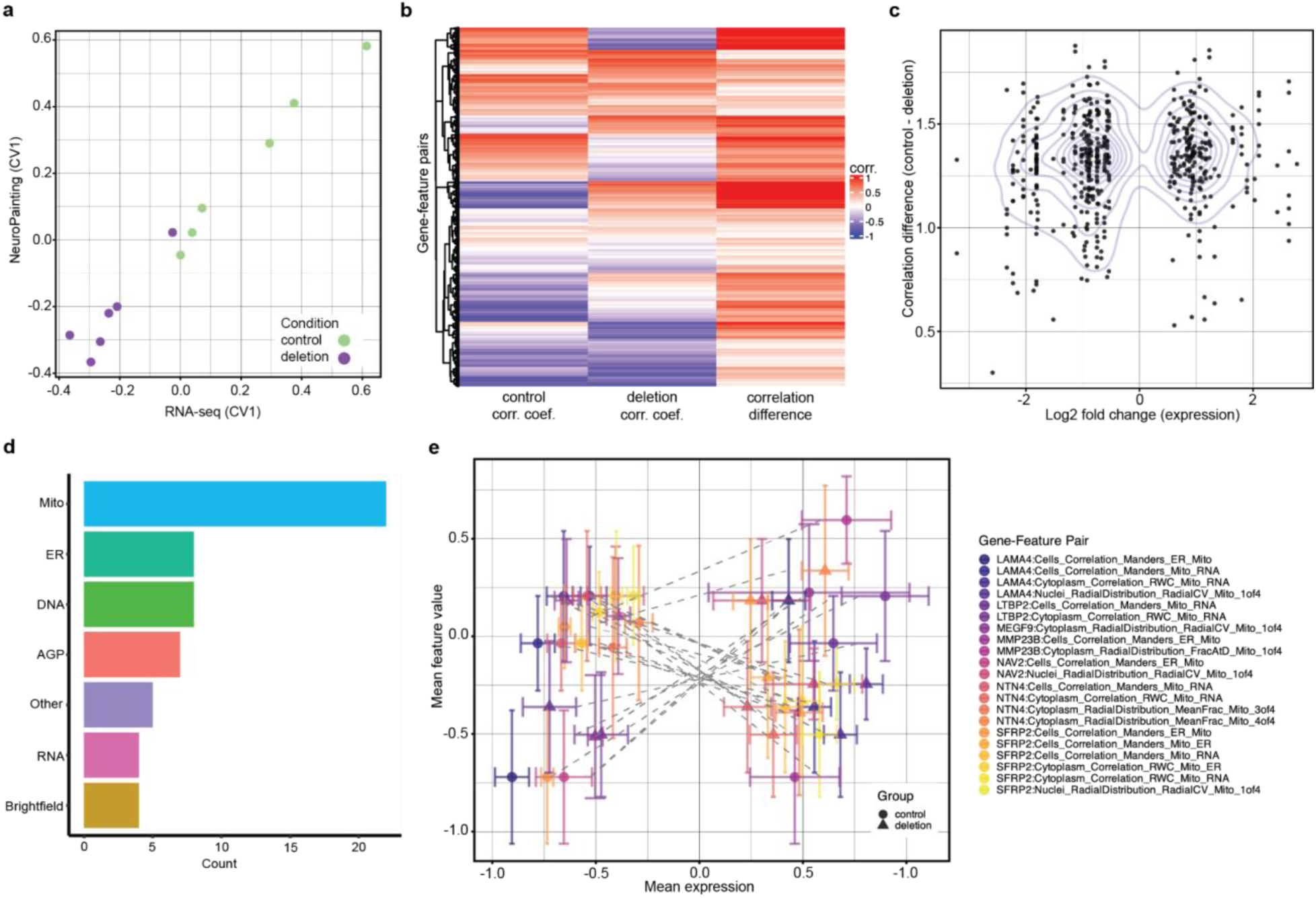
Integration of RNA-seq and NeuroPainting data links cell adhesion gene expression to mitochondrial morphology. **a.** Scatter plot of CCA components for RNA-seq and NeuroPainting profiles, colored by condition (control = green, 22q11.2 = purple). **b.** Heatmap of correlation coefficients between gene expression and morphology feature values for top 100 genes and top 100 features from CCA loadings. Correlation difference is measured as the absolute value between control and deletion coefficients. **c.** Scatter plot of absolute differences in correlation coefficients against fold change coefficients between control and 22q11.2 deletion astrocytes for gene-features pairs. **d**. Barplot for morphology feature categories for cell adhesion gene-feature pairs. **e**. Scatter plot comparing mean gene expression and morphology values for cell adhesion gene-mitochondrial feature pairs. Each data point represents a specific gene-feature pair, with different colors used to distinguish pairs. Error bars represent the standard error of the mean for both expression (horizontal) and morphology (vertical). Dashed lines connect the control and deletion points for each gene-feature pair.

Then we more thoroughly examined the specific relationships between these 100 genes and 100 features to identify relationships between these pairs which were altered by the presence of the 22q11.2 deletion. We computed pairwise correlations for each gene-feature pair, separately for control and deletion conditions, and identified 530 gene-feature pairs with significantly different correlations (R^2^) between control and deletion samples (**Fig. 5b**, p-value < 0.05).

We were particularly interested in gene-features pairs whose correlations displayed dramatic changes. For example, whether there were pairs with strong positive correlations in control samples, but in the presence of the deletion showed strong negative correlations. Changes in correlation were quantified as the difference in correlation coefficients between conditions. We observed many gene-feature correlations that were dramatically different between the control and 22q11.2 deletion astrocytes. Notably, features related to RadialDistribution and Granularity were disproportionately affected (chi-square test, χ² = 23.669, df = 5, p = 0.00025). This suggested that many of our gene expression changes were associated with these feature categories.

To investigate whether these changes in gene-morphology correlations were driven by underlying alterations in gene expression, we examined the relationship between log2 fold change (log2FC) in gene expression and the observed differences in correlation coefficients using Spearman’s rank correlation test. We found a significant negative correlation (rho = - 0.364, p-value < 1e-05) suggesting that genes with larger changes in expression (either upregulated or downregulated) tended to exhibit more substantial changes in their correlations with morphological features in the deletion samples (**Fig. 5c**).

Next, we wanted to explore the biological context of these relationships. To do so, we conducted a pre-ranked GSEA including the genes from our significant gene-feature pairings. We identified two pathways which were significantly enriched in our list (FDR-adjusted p < 0.05). The pathway “glial cell apoptotic process” (GO:0034349) was enriched, driven by three key genes: *PRKCH, AKAP12*, and *TNFRSF21*. Enrichment of this pathway suggests the 22q11.2 deletion may be rendering astrocytes more vulnerable to apoptosis. Additionally, the “collagen-containing extracellular matrix” pathway (GO:0062023) included nine genes (*LTBP2, LAMA4, SBSPON, NAV2, COLEC12, NTN4, MMP23B, MEGF9, and SFRP2*), many of which are implicated in cell adhesion processes ^50–53^. This finding corroborates our earlier results that 22q11.2 deletion may alter genes associated with cell adhesion and membrane structure.

Then we sought to identify specific morphological changes which might be linked to the genes in our GSEA. When we examine the features associated with the extracellular matrix pathway, we observe a strong enrichment of mitochondrial associated features. (**Fig. 5d**, chi-square test, χ² = 28.655, df = 6, p-value = 7.069e-05), suggesting that changes in the expression of these nine genes elicited a specific disruption of mitochondrial function and organization relative to all other feature types. When examining individual gene-feature pairs, we found pronounced differences in correlation coefficients between control and 22q11.2 deletion samples for cell adhesion genes and mitochondrial features (**Fig. 5e**).

## Discussion

This study presents the development and application of NeuroPainting, a high-dimensional phenotyping assay adapted from Cell Painting, to identify morphological phenotypes in iPSC-derived neural cell types, including stem cells, astrocytes, neural progenitors, and neurons. Our results demonstrate the assay’s robustness and utility to uncover novel insights into neuropsychiatric disorders, particularly those linked to the 22q11.2 deletion syndrome, a major genetic risk factor for schizophrenia. Through this work, we sought to address several key challenges in cellular and molecular psychiatric research, including the need for disease-relevant cellular models and the ability to capture complex, cell-type-specific phenotypes.

NeuroPainting facilitates the implementation of scalable genetic and chemical screening assays, helping to catalyze target and therapeutic discovery in neuropsychiatric research. The classification of over 4,000 cellular traits into well-defined categories such as size, shape, and texture across different cellular compartments allows for high-resolution analysis of neural cell morphology. Moreover, the extensive public dataset we created serves as a gold standard for the research community, enabling others to compare and validate their results, thus setting a new benchmark for phenotypic screening in neuropsychiatric research.

We showed that NeuroPainting can successfully distinguish cell types based on morphological profiles. It can thus be leveraged to explore possible alterations in cell state (for instance following genetic or pharmacological perturbations), and whether genetic mutations or drug treatments impact differentiation potential. In approaches where multiple cell types are present in the population that is being screened, this tool could also reveal shifts in cell type composition based on cellular morphology.

The application of NeuroPainting to iPSC-derived neural cells from individuals with the 22q11.2 deletion syndrome revealed diverse, cell-type-specific morphological signatures associated with this genetic mutation. Notably, the deletion’s effects were most pronounced in stem cells and astrocytes, with many morphological features significantly altered in these cell types. The morphological signatures identified in astrocytes, particularly those related to the radial distribution and localization of mitochondria, ER, and AGP, point to new biological mechanisms that may underlie the neuropsychiatric symptoms associated with the 22q11.2 deletion. The altered relationship between the ER and mitochondria, as indicated by the significant differences in Cytoplasm_Correlation_RWC_ER_Mito, could represent altered mitochondria-associated endoplasmic reticulum membranes (MAMs), which play a crucial role in diverse cellular functions, including metabolic regulation, signal transduction, autophagy, and apoptosis ^54^.

We provided the first investigation of transcriptional dysregulation in iPSC-derived astrocytes with a 22q11.2 deletion, building on our past findings of 22q11.2 deletion associated transcriptional changes in iPSC-derived neural progenitor cells and excitatory neurons ^13^. Our findings suggest that the deletion reduces the expression of many neurodevelopmental gene sets, including many genes that play a role in cell adhesion and migration. This result was interesting as recent studies have implicated altered expression of astrocytic cell adhesion genes in schizophrenia ^40,55^. This is the first indication that the Synaptic Neuron Astrocyte Program (SNAP) is reduced in (living) human astrocytes with 22q11.2 deletion. Previously, decreased SNAP expression had only been observed in astrocytes from post-mortem brains of individuals with idiopathic schizophrenia ^39^. Additionally, the integration of NeuroPainting and transcriptomic data identifies cell morphological phenotypes downstream of these changes in gene expression that are associated with mitochondrial morphology. These results validate many previous reports of mitochondrial dysfunction in 22q11.2 syndrome and neuropsychiatric disorders more broadly ^56,57^. Our hope is that the broader context of linking gene expression to these mitochondrial phenotypes may provide a novel inroad into understanding these pathological mechanisms. In this instance, we defined a relationship between what is emerging as a robust cellular program altered in persons with schizophrenia with a morphological trait that is screenable *in vitro*.

Our study has limitations that should be addressed in future research. While we reliably identified morphological signatures in stem cells, progenitors, and astrocytes, we encountered greater difficulty with neurons. This could be due to the complex morphology of neurons, resulting in perhaps noisier data with higher variability between samples. Larger sample sizes may help mitigate this challenge. It is also possible that the effects of the 22q11.2 deletion may produce more subtle phenotypes in neurons compared to the other cell types studied. Additionally, although our approach captures thousands of morphological features, it does not include specific phenotypes commonly analyzed in neuroscience, such as synaptic phenotypes. Integrating NeuroPainting with synapse-specific markers could enable more targeted analysis, combining broad-scale morphological profiling with traditional, well-established measurements familiar to the neuroscience field.

This study demonstrates the power of NeuroPainting to uncover novel morphological phenotypes associated with neuropsychiatric conditions. The cell-type specificity of these phenotypes, coupled with the integration of gene expression data, provides a deeper understanding of the underlying mechanisms of 22q11.2 deletion syndrome. This was the first study to show that the 22q11.2 deletion reduces the expression of cell adhesion genes, which mirrors recent findings from post-mortem brain studies of schizophrenia. Excitingly, we link these changes to specific morphological traits, including altered mitochondrial phenotypes. These relationships may provide new insights into ways that transcriptional changes in the human brain may impact cell function. Future studies could expand the application of NeuroPainting to additional neural cell types and could explore phenotypes associated with other genetic mutations or chemical perturbations. Ultimately, this approach holds promise for the development of new therapeutic strategies targeting the cellular and molecular bases of neuropsychiatric disorders.

## Materials and Methods

### Human pluripotent stem cell (hPSC) lines cohort and derivation

We used a cohort of cell lines we previously described (Nehme et al, 2022). Briefly, we assembled a scaled discovery sample set through highly collaborative, multi-institutional efforts with the Stanley Center Stem Cell Resource (Broad Institute), the Swedish Schizophrenia Cohort (Karolinska Institute), the Northern Finnish Intellectual Disability Cohort (NFID), Umea University, Massachusetts General Hospital (MGH), McLean Hospital, and GTEx.

### iPSC culture

Human iPSCs were maintained on plates coated with Geltrex (life technologies, A1413301) in StemFlex media (Gibco, A3349401) and passaged with accutase (Gibco, A11105). All cell cultures were maintained at 37 °C, 5% CO2.

### Neuronal progenitor and excitatory neuron differentiation

iPSCs were differentiated into cortical glutamatergic neurons as previously described (Nehme et al, 2018). On D0 iPSCs were passaged and replated at a density of 1M cells per will in a 6-well plate. On D1 iPSCs were differentiated in Neural Induction Medium (NIM) [500 mL DMEM/F12 (1:1) (Gibco, Cat # 11320-033), 5 mL Glutamax (Gibco, Cat # 35050-061), 7.5 mL Dextrose (20%, SIGMA, Cat # 1181302), 5 mL N2 supplement (Invitrogen, Cat # 17502048)] supplemented with SB431542 (10 μM, Stemcell Technologies, Cat # 72234), XAV939 (2 μM, Stemcell Technologies, Cat # 72672) and LDN-193189 (100 nM, Stemcell Technologies, Cat # 72147) along with doxycycline hyclate (2 μg.mL-1, Sigma, Cat # D9891). Media was replaced on D2 and D3 with the same media used for D1 with the addition of Zeocin (1 ug/ml). On D4, cells are co-cultured with mouse glia to promote neuronal maturation and synaptic connectivity (Eroglu et al, 2010, Pfrieger et al, 2009) (TransnetYX, Cat # C57EASTWB). From D4 onward, cells are maintained in Neurobasal media (500 mL Neurobasal [Gibco, 21103-049], 5 mL Glutamax [Gibco, 35050-061], 7.5 mL Dextrose [20%, SIGMA, Cat # 1181302], 2.5 mL MEM-NEAA [Invitrogen, Cat # 11140050]) supplemented with B27 (50X, Gibco, 17504-044), BDNF, CTNF, GDNF (10 ng.mL–1, R&D Systems 248-BD/CF, 257-NT/CF, and 212-GD/CF) and doxycycline hyclate (2 µg.mL–1, Sigma, D9891). From day 4 to day 5, Neurobasal media was complemented with the antiproliferative agent floxuridine (10 µg.mL–1, Sigma-Aldrich, Cat # F0503-100MG). To capture neuronal progenitor cells, we harvest D4 samples prior to adding them to the mouse glia.

### Astrocyte differentiation

iPSCs were differentiated into astrocytes as previously described (Berryer et al, 2023). On D0 iPSCs were passaged and replated at a density of 1M cells per will in a 6-well plate. On day 1, iPSCs were differentiated in N2 medium [500 mL DMEM/F12 (1:1) (Gibco, Cat # 11320-033), 5 mL Glutamax (Gibco, Cat # 35050-061), 7.5 mL Dextrose (20%, SIGMA, Cat # 1181302), 5 mL N2 supplement B (StemCell Technologies, Cat # 07156)] supplemented with SB431542 (10 μM, Stemcell Technologies, Cat # 72234), XAV939 (2 μM, Stemcell Technologies, Cat # 72672) and LDN-193189 (100 nM, Stemcell Technologies, Cat # 72147) along with doxycycline hyclate (2 μg.mL-1, Sigma, Cat # D9891). Media was replaced on D2 with the same media used for D1 with the addition of Zeocin (1 ug/ml). Starting on day 3 human induced neural progenitor-like cells were harvested with Accutase (Innovative Cell Technology, Inc., Cat # AT104-500) and re-plated at 15,000 cells cm-2 in Astrocyte Medium (ScienCell, Cat # 1801) with Y27632 (5 mM) on geltrex coated plates. Cells were maintained for 30 days in Astrocyte Medium (ScienCell, Cat # 1801).

### Cell seeding and staining

For each batch of imaging, cells were detached from 6-well NUNC plates using Accutase (StemcellTech; cat#07920) for generating single-cell suspensions. Following detachment, cells were centrifuged at 1000 rpm x 5:00 and re-suspended in the appropriate medium for each individual cell type. After each cell line was counted to determine cell solution concentration and viability, the desired cell solution volume was aliquoted into a 96-deep well low attachment plate following a specific plate map to ensure that wells from any given cell line were not predominantly on the edge wells or too close together. To disperse a high number of cell lines across a 384-well plate in a semi-random fashion, we optimized the use of an Agilent Bravo liquid handling device. Here, using an 8-channel head, cell solutions were transferred from the 96-well low attachment plate and distributed into a geltrex-coated Perkin Elmer Cell Carrier 384-well plate at the determined density per well. Each cell line was plated into 8 distinct wells on the final screening plate in four-well quadrants. The ideal seeding densities were determined through pilot experiments and evaluated based on visual assessment of the cells. The conditions used in this study are outlined in **Fig. 1b**.

### NeuroPainting and imaging

Cells were stained and imaged with minor adaptations to procedures described previously ^17^. Post seeding in 384-well plates, cells were treated for 30min with 0.5 uM MitoTracker Deep Red FM - Special Packaging (Thermo Fisher cat#: M22426) dye at 37oC. Following the MitoTracker treatment, cells were fixed with 16% paraformaldehyde diluted to a final concentration of 4% (Thermo Fisher cat#: 043368.9M) for 20min in the dark at RT. After three washes with 1X HBSS cells were permeabilized and stained using a solution of 1X HBSS (Thermo Fisher cat#: 14175095), 0.1% Triton-X-100 (Sigma Aldrich cat#: X100-5ML), 1% Bovine Serum Albumin, 8.25 nM Alexa Fluor 568 Phalloidin (Thermo Fisher cat#: A12380), 0.005mg/ml Concanavalin A, Alexa Fluor 488 Conjugate (Thermo Fisher cat#: C11252), 1ug/ml Hoechst 33342, Trihydrochloride, Trihydrate (Thermo Fisher cat#: H3570), 6uM SYTO 14 Green Fluorescent Nucleic Acid Stain (Thermo Fisher cat#: S7576), and 1.5ug/ml Wheat Germ Agglutinin, Alexa Fluor 555 Conjugate (Thermo Fisher cat#: W32464) for 1 h at RT in the dark. Following the staining, plates were washed 3X with 1X HBSS and sealed until imaging. Cell Painted plates were imaged on a Perkin Elmer Phenix Automated Microscope under a standardized protocol.

### Quantification of cellular morphology traits and their quality control

The segmentation of individual cells in the image into their cellular compartments (whole cell, cytoplasm and nuclei) and subsequent quantification of morphology traits for each cellular compartment was done using CellProfiler 4.2.4 (Stirling et al, 2021; pipelines are available at https://github.com/broadinstitute/NeuroPainting). Subsequently, cells missing measurement for more than 5% of traits were removed. Morphology traits a priori known to be problematic, not measured across all cells or non-variable across cells were removed using Caret v6.0-86 package. QC-ed cells were then segregated into two groups based on the number of neighbors: isolated cells having no neighbors and colony cells having one or more neighbors. Individual morphology traits were then summarized to well level measurement by averaging them across all cells per well, resulting in a well by trait matrix. Following this, each morphology trait was gaussianized across all 4 plates using inverse normal transformation (INT) method.

### scRNA-sequencing and donor assignment

For single-cell analyses, cells were harvested and prepared using 10X Chromium Single Cell 3’ Reagents v3 and sequenced on an Illumina NovaSeq 6000 with an S2 flow cell, generating paired-end reads of 2 x 100 bp. Raw sequencing data were aligned and processed following the Drop-seq workflow. Human reads were aligned to the GRCh38 reference genome and filtered for high-quality mapped reads (mapping quality ≥ 10). To determine the donor identity of each droplet, variants were filtered through multiple quality controls, ensuring only high-confidence A/T or G/C sites were included in the VCF files. Once the single-cell libraries and VCF reference files were filtered and quality-checked, the Dropulation algorithm was applied. This algorithm analyzes each droplet (or cell) independently, assigning a probability score to each variant site based on the observed versus expected allele. Donor identity is determined by computing the diploid likelihood at each UMI, summed across all sites, to identify the most likely donor for each droplet. To ensure consistency in differential expression testing downstream, we remove any cells which are clustered separately by individual donors.

### Random forest classification for cell types

A random forest classifier was used to classify cell types based on morphological profiles from control samples. The data was split into 70% training and 30% testing sets, and a 5-fold cross-validation was employed to optimize hyperparameters. The number of features considered at each split (mtry) was tuned using a grid search with values ranging from 2 to 721. The final model was built using the best-performing mtry, with 500 trees, a maximum of 60 terminal nodes, and a minimum node size of 5. Model performance was evaluated on the test set using a confusion matrix, accuracy, sensitivity, specificity, and Kappa statistics. Receiver Operating Characteristic - Area Under the Curve (ROC-AUC) scores were computed for each cell type, and predicted probabilities were visualized using boxplots. All analyses were performed in R (version 4.0.2) with the randomForest, caret, and pROC packages, and a random seed of 42 to ensure reproducibility.

### Differential feature analysis

To identify differential features between control and deletion conditions across various cell types, we conducted a Wilcoxon rank-sum test using the well-level NeuroPainting profiles. The dataset was grouped by *Metadata_cell_type* to ensure comparisons were made within each cell type. For each group, a pairwise Wilcoxon rank-sum test was performed to compare control and deletion conditions for each numeric feature, with p-values recorded. To address multiple comparisons, we applied a False Discovery Rate (FDR) correction to the p-values within each cell type using the Benjamini-Hochberg (BH) method.

### Differential gene expression

Differential gene expression analysis was conducted using DESeq2 (v1.34.0), a statistical package designed for analyzing count data from RNA sequencing experiments. Pseudo-bulk RNA expression profiles for each cell line were generated by summing the individual cell counts across each sequencing reaction, and these counts were used as input for DESeq2. The data were pre-processed to ensure quality control, including filtering out lowly expressed genes that had fewer than 10 reads in at least 50% of the samples.Normalization of the count data was performed using the DESeq2 default method, which estimates size factors for each sample to account for differences in sequencing depth. These size factors were used to normalize the read counts across samples, ensuring comparability. The DESeq2 model was fitted to the data using the negative binomial distribution to estimate the dispersion for each gene. Differential expression was determined by comparing the expression levels between the conditions of interest. For this study, the primary comparison was between 22q11.2 deletion and control, with the design formula in DESeq2 specified as *∼ condition*, where *condition* represents the categorical variable of interest. DESeq2 calculates the log2 fold change (log2FC) for each gene, representing the ratio of gene expression between the two conditions. Statistical significance was assessed using the Wald test, and p-values were adjusted for multiple testing using the Benjamini-Hochberg procedure to control the false discovery rate (FDR). Genes with an adjusted p-value (padj) of less than 0.05 were considered significantly differentially expressed. A total of 835 genes were identified as differentially expressed between the conditions, with thresholds of log2FC > |1| and padj < 0.05.

### Canonical Correlation Analysis (CCA)

We performed Canonical Correlation Analysis (CCA) to investigate the relationship between RNA sequencing (RNA-seq) gene expression data and morphological features (NeuroPainting). We used inverse normalized transformed data including the DEGs (1358) and significant features (536). To reduce noise and multicollinearity, we first applied Principal Component Analysis (PCA) to both datasets, retaining the top 5 principal components (PCs) from each. For RNA-seq data, PCA was performed on z-score expression values, while for morphological features, PCA was applied to z-scored feature values. The CCA was then conducted using the cancor() function in R, with the top 5 PCs from each dataset serving as input. We extracted the loadings from the first canonical variable (CV1) to identify the top contributing genes and morphological features for subsequent analysis, providing insight into the shared variance between the two datasets.

### Morphology and Gene Expression Correlation Analysis

We performed a fixed-effects linear regression analysis to investigate the relationship between gene expression and morphological features in astrocytes with and without the 22q11.2 deletion. The model included gene expression as the predictor and various morphological traits as the response variable, with genotype as an interaction term to test for differential effects between control and deletion groups. Specifically, we fit the following model for each gene-feature pair:

Morphology ∼ Expression * Group

The interaction term was used to assess whether the correlation between gene expression and morphology varied between the control and deletion groups. p-values for the interaction term were calculated, and significant gene-feature pairs were identified. Coefficients for expression and the interaction term were extracted to assess the strength and direction of the gene-morphology relationships across groups.

### Gene Set Enrichment Analysis (GSEA)

We performed a pre-ranked gene set enrichment analysis (GSEA) using the clusterProfiler package in R (v4.6.0). We employed the gseGO() function with the Gene Ontology (GO) database, querying both the Biological Process (BP) and Cellular Component (CC) ontologies. Gene symbols were mapped to Entrez IDs using the bitr() function and the org.Hs.eg.db annotation package (v3.16.0). To ensure accurate mapping, we filtered out genes without valid Entrez identifiers, resulting in a final gene list for enrichment analysis. The GSEA was performed using default parameters, with the following settings: minimum gene set size of 10, maximum gene set size of 500, and permutation type set to gene labels. Significance thresholds were determined using the Benjamini-Hochberg procedure to control the False Discovery Rate (FDR). Pathways with an adjusted p-value (FDR) < 0.05 were considered statistically significant. Enrichment scores (ES) and normalized enrichment scores (NES) were used to rank the significance of enriched pathways, and results were visualized using the enrichplot package.

### Identification of Significant Gene-Feature Pairs

We calculated Pearson correlation coefficients for gene expression and morphology for both control and deletion groups, deriving the absolute difference between these two correlation coefficients for each pair. Significant gene-feature pairs were defined as those with an interaction p-value < 0.01 and a correlation difference > 1.0. Fisher’s Z-test was applied to evaluate whether the correlation differences were statistically significant.

### Enrichment Analysis for Feature Categories and Cellular Compartments

To assess whether certain morphological features or cellular compartments were overrepresented among the significant gene-feature pairs, we conducted a Chi-squared independence test. We grouped features by their respective categories (e.g., Granularity, Texture, Correlation) and compartments (e.g., Nuclei, Cytoplasm, Cells) and compared their distribution in our set of gene-pairs to their overall distribution amongst the full feature set.

–---

## Data availability

The raw image data will be available in the Cell Painting Gallery on the Registry of Open Data on AWS (https://registry.opendata.aws/cellpainting-gallery/) as dataset cpg0038-tegtmeyer-neuropaintingat no cost and no need for registration. Sequencing data, including gene expression matrices will be made available on the Broad Institute Single-cell Portal.

## Code Availability

Source code to reproduce and build upon the presented results is available at https://github.com/broadinstitute/NeuroPainting.

## Acknowledgments

With thanks Kris Dickson for reviewing and editing the manuscript, and members of the Nehme lab and the Carpenter-Singh lab for helpful comments and discussion.

## Contributions

R.N. and S.S. conceived the work. M.T. and R.N. wrote the manuscript with input from all authors. M.T. performed the experiments with help from D.L., K.B. and D.H. M.T, P.V.R., C.T.-C., B.A.C., S.S., G.W., M.H. processed and analyzed Cell Painting data. R.N., A.E.C., and S.S. supervised the work and analyses. All authors edited and approved the final manuscript.

## Supplementary Figures

**Supplementary Figure 1.**
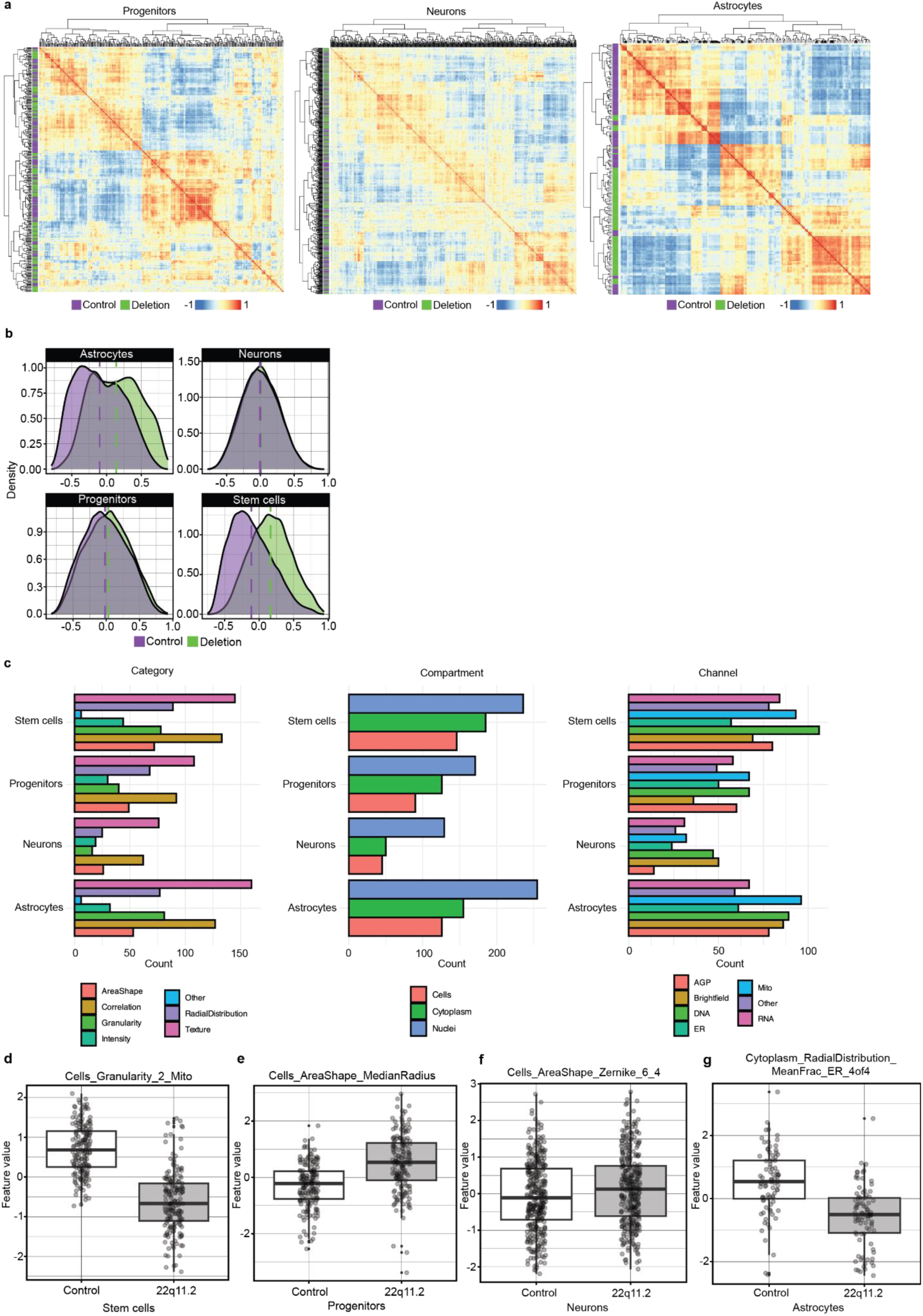
a. Heatmaps for pairwise-pearson correlations between NeuroPainting profiles across progenitors, neurons, and astrocytes. Colored annotation bar denotes genotypes for control (purple) and 22q11.2 deletion samples (green). b. Distribution of pairwise Pearson correlations for control-control (green) and control-deletion (purple) samples for each of the four cell types: stem, progenitor, neuron, and astrocyte. Vertical error bars indicate mean correlations. c. Barplot of the number of significant features by cell type (Wilcoxon Rank-Sum test, FDR < 0.05). c. Barplot for significant features (FDR < 0.05) between control and 22q11.2 deletion samples across cell types. d-g. Boxplots for representative significant features in stem cells, progenitors, neurons, and astrocytes, presented like Fig. 2f.

**Supplementary Figure 2.**
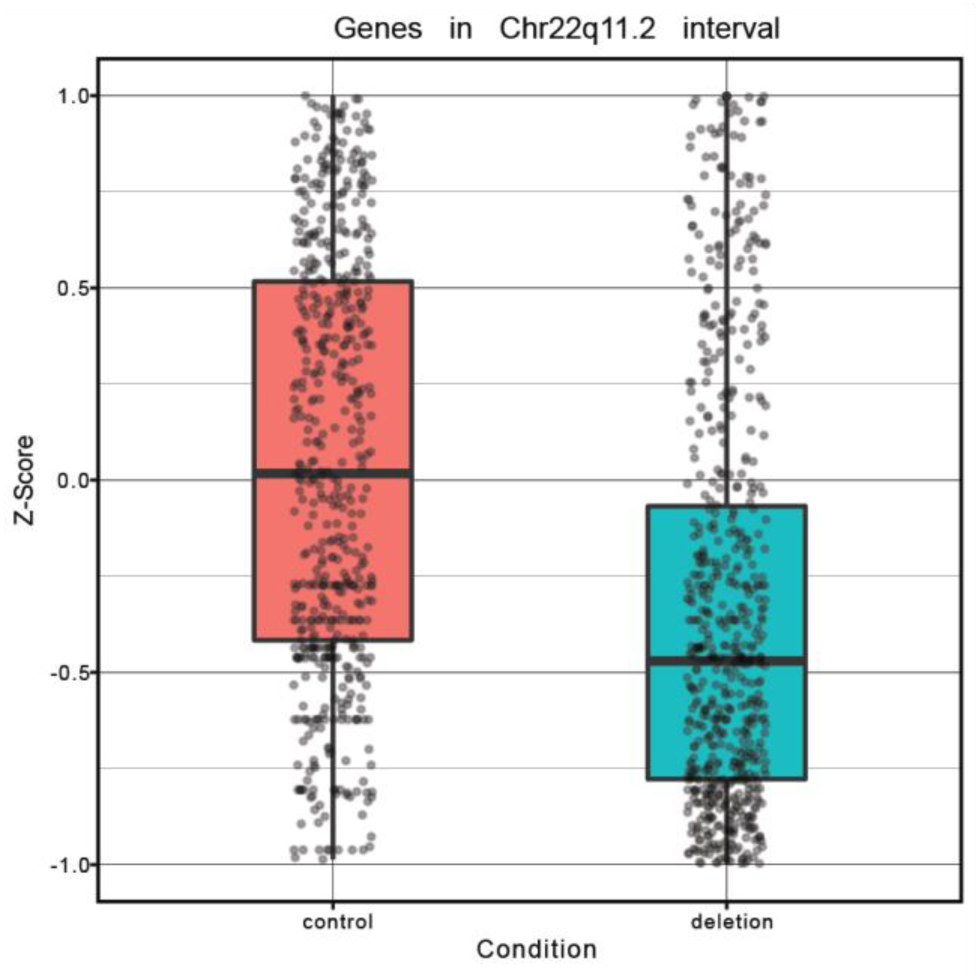
Z-scored expression of genes within the 22q11.2 interval. Data are presented in a Tukey-style boxplot with the median (Q2) and the first and the second quartiles (Q2, Q3) and error bars defined by the last data point within ±1.5-times the interquartile range.

